# CELL-E: A Text-To-Image Transformer for Protein Localization Prediction

**DOI:** 10.1101/2022.05.27.493774

**Authors:** Emaad Khwaja, Yun S. Song, Bo Huang

## Abstract

Accurately predicting cellular activities of proteins based on their primary amino acid sequences would greatly improve our understanding of the proteome. In this paper, we present CELL-E, a text-to-image transformer architecture that generates a 2D probability density map of protein distribution within cells. Given a amino acid sequence and a reference image for cell or nucleus morphology, CELL-E offers a more direct representation of protein localization, as opposed to previous *in silico* methods that rely on pre-defined, discrete class annotations of protein localization to subcellular compartments.

## 1 Introduction

In recent years, advancements in sequencing technologies have allowed for the comprehensive cataloging of proteins and their amino acid sequences across a wide range of organisms [1]. Despite this progress, the exact functions and cellular dynamics of many proteins remain unclear. In order to gain a deeper understanding of these proteins, researchers have sought ways to predict their properties, including structure, interactions, subcellular localization, and trafficking patterns, from their amino acid sequences. This type of computational analysis has the potential to shed light on the “dark matters” of the proteome and enable large-scale screening before expensive experimental validation. These tools have numerous applications in biomedical research, such as drug design and therapeutic target discovery [2].

In this study, our focus is on predicting subcellular localization of proteins from their amino acid sequences, which serves as the spatial context for their cellular functions. The localization of a protein to a specific subcellular compartment can be driven by either active transport or passive diffusion in conjunction with specific protein-protein interactions, often involving localization “signals” in the amino acid sequence [3–5]. In many cases, however, the exact mechanisms for sequence recognition and trafficking are not yet fully understood [6]. For example, there is ongoing debate about the mechanism behind the import of proteins via the nuclear localization sequence (NLS) [7]. Given these challenges, machine learning utilizing existing knowledge of protein localization has become a particularly useful tool.

Although computational prediction of protein subcellular localization from primary amino acid sequences is an active area of research, most works train the model with class annotation of subcellular compartments (e.g. nucleus, plasma membrane, endoplasmic reticulum, etc.) which are available from databases such as UniProt [8]. This approach has two major limitations. First, many proteins are present in different and variable amounts across multiple subcellular compartments. Second, protein localization could be highly heterogeneous and dynamic depending on the cell type and cell state (including cell cycle state). Neither of these two aspects have been captured by existing discrete class annotations. Consequently, machine-learning-based protein localization prediction still have limited applications [9]. Furthermore, to assist mechanistic discoveries, it is highly desirable for the machine learning models to be explainable.

To investigate the relationship between sequence and subcellular localization, we present CELL-E, a text-to-image based transformer which predicts the probability of protein localization on a per-pixel level from a given amino acid sequence and a conditional reference image for the cell morphology and location (Fig. 1). It relies on transfer learning via amino acid encodings from a pretrained protein language model, and two quantized image encoders trained from a live-cell imaging dataset. By generating a two-dimensional probability density function (2D PDF) atop the reference image, CELL-E naturally accounts for multi-compartment localization and the cell type/state information implicitly encoded by the cell morphology. We demonstrate the capability of CELL-E to predict localization of proteins, identify changes in localization from mutations, and uncover sequence features correlated with the specification of subcellular protein localization.

**Fig. 1.**
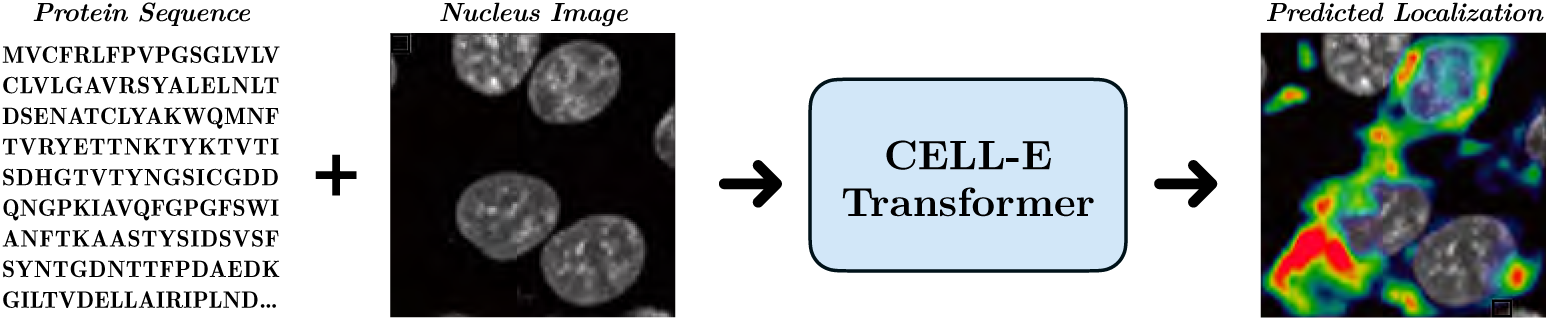
Given an input of amino acids and a reference nucleus image, CELL-E makes a prediction of protein localization with respect to the nucleus as a 2D probability density function, shown as heatmap, with color indicating relative confidence for each pixel.

**Fig. 2.**
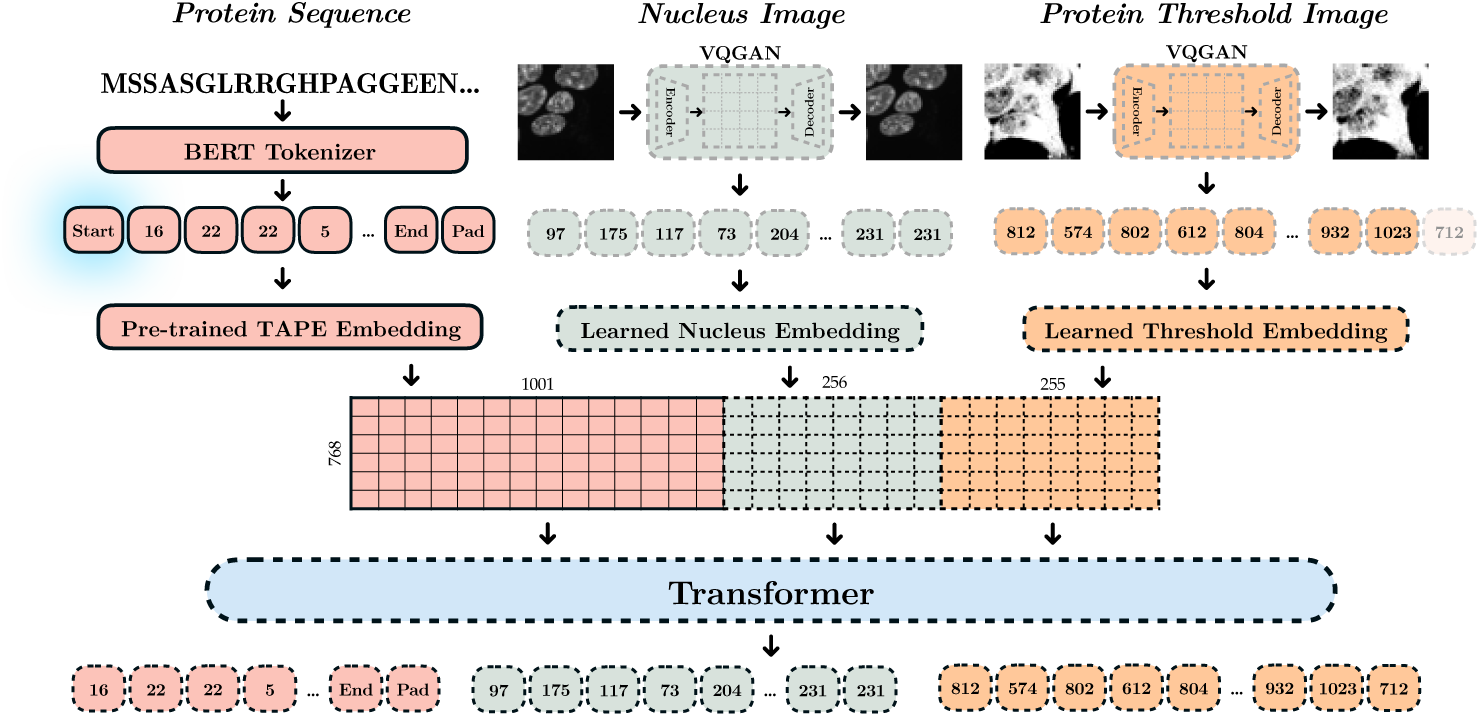
Graphical depiction of CELL-E. Solid lines correspond to pre-trained components. Gray dashed lines are learned in Phase 1 and 2 (Reference Image and Protein Threshold VQGANs). Black dashed lines correspond to components learned in Phase 3. A start token is prepended to the sequence and the final protein image token is removed. The amino acid sequence embedding from the model is preserved, and embedding spaces for the image tokens are cast in the same depth and concatenated with the amino acid sequence embedding. The transformer is tasked with reproducing the original sequence of tokens (e.g. the input sequence with start token shifted to the right one position).

## 2 Results

### 2.1 The CELL-E Model

CELL-E is inspired by the text-to-natural-image generation model of DALL-E [10] (See Section S.2.1) for a review of relevant work). Similar to DALL-E, our model autoregressively learns text and image tokens as a single stream of data. While the goal of general text-to-image models is to produce images with high perceptual strength, they do not necessarily aim for quantitative accuracy [10–12]. Therefore, CELL-E was designed with the following considerations:

1. **Transfer learning.** Training CELL-E requires a library of cellular images and corresponding morphological reference images for a large number of proteins. For this purpose, we utilized the recently established OpenCell library[13], which contains a library of 1,311 CRISPR-edited HEK293T human cell lines, each having one target protein fluorescently tagged and imaged by confocal microscopy with accompanying DNA staining as the reference for nuclei morphology. The high image quality and consistency makes OpenCell a good choice as the training and validation dataset (See Section S.3.1 for more information). Still, data availability in this domain remains a large obstacle. For example, DALL-E was trained on 250 million text-images pairs [10], whereas even the largest publicly available dataset with annotated protein images in human cells, Human Protein Atlas (HPA), only contains 12,003 unique proteins with just 82,000 images [14]. We found that utilizing transfer learning by incorporating a frozen embeddings from a language model pre-trained over thirty-one million protein domains from Pfam [15] as the input embedding for the amino acid text sequence. This approach reduces the number of learned paramaters, thereby alleviating the burden for CELL-E to also learn the amino acid sequence space. This allows training to be concentrated on the relationship between sequence and image tokens. We evaluated multiple protein language models (see Supplementary Information and Table S.1) and eventually chose the BERT-based model from Rao et. al. [16], which we refer to as the TAPE model, for subsequent work.

2. **Morphological reference.** In our initial efforts, we found that a transformer using just the amino acid tokens and image tokens is capable of generating cell-like images from the amino acid sequence alone (Fig. S.3). However, quantifying protein localization information in the generated images is challenging. Furthermore, an estimation of a single snapshot of protein localization is not necessarily a quantifiable indication of global behavior. Therefore, in addition to amino acid tokens and protein image tokens, we utilize 3 separate embedding spaces that also include tokens representing the overall cell morphology from a reference image. The references image provides the model with information regarding the localization of subcellular structures and compartments. Moreover, cell morphology implicitly provides the cell type and cell state context for CELL-E predictions.

3. **Image model.** Instead of the Vector Quantized Varational Autoencoder (VQVAE) previously used to analyze OpenCell imaging data [17], we chose to use Vector Quantized Generative Adversarial Network (VQGAN) [18] which produces images with comparatively higher spatial frequency. To simplify the task of the protein image VQGAN, we let it predict per-pixel binary representations of protein localization (i.e. a thresholded image). This allows us to use the marginal probabilities predicted for each image token from CELL-E to create a weighted sum on the image tokens. This latent space linear combination is then used to generate a continuous 2D probability density function of protein localization, which resembles a gray-scale image (Fig. 7). We note that the same model can also be trained to output gray-scale images directly (See Supplementary Notes S.2.2 and Fig. S.4).

### 2.2 Performance Evaluation

Fig. 3 and Fig. S.2 shows the CELL-E predictions for several proteins in the validation dataset. High similarities can be seen between the predictions and the ground truth. Even though the reference images only depict the nuclei, which is a limitation of the OpenCell training data, CELL-E can reasonably paint the shape of the cell for cytoplasmic proteins. Interestingly, the case of Mitogen-Activated Protein Kinase 9 (MAPK9) contains a cell in metaphase. CELL-E correctly predicts the round shape of its distribution around the mitotic chromosomes instead of the more expanded distribution for the adjacent interphase cell. This result suggests that CELL-E can indeed capture cell state information from the morphological reference images.

**Fig. 3.**
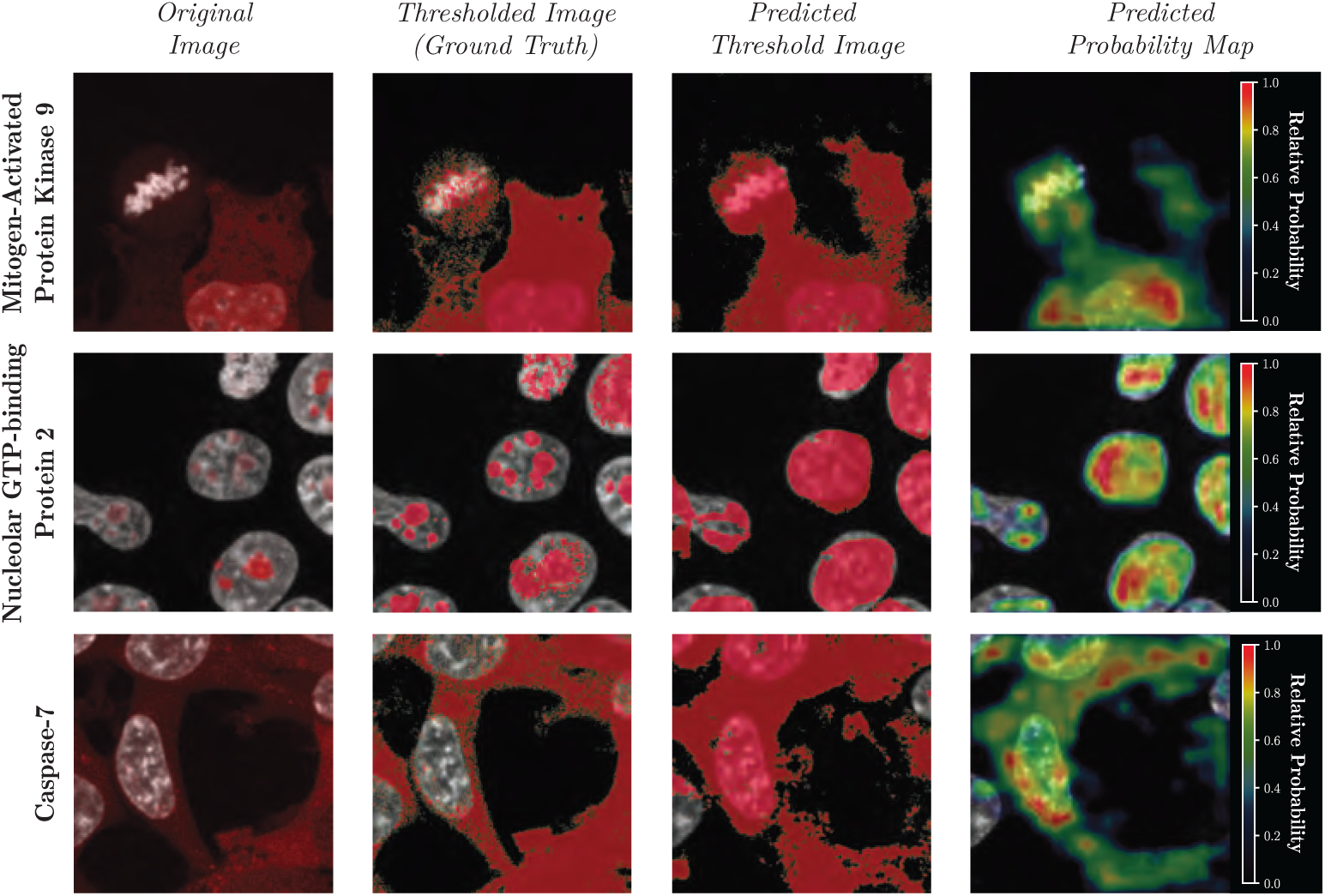
Prediction results of several types of proteins from the validation set, unseen to the model during training. The nucleus channel is depicted in grayscale, and the protein channel is shown as an overlay in red (Fig. S.1 for clarification). The thresholded image (Column 2) is designated “Ground Truth” because those are the types of images exposed to the model during training. The predicted probability map is obtained from a weighted sum of potential image patches and normalized to 1.

**Fig. 4.**
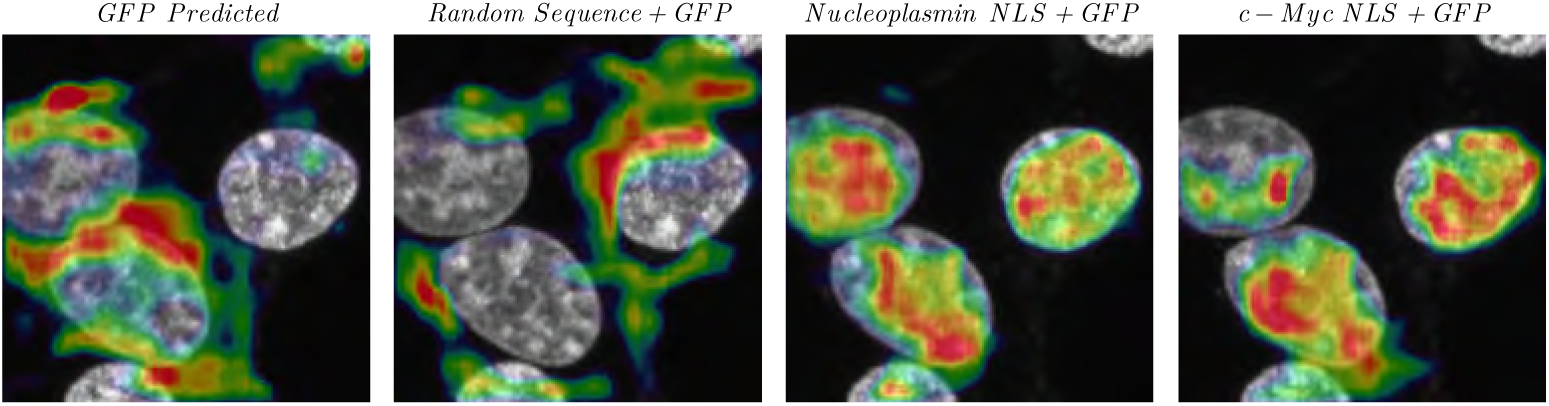
Predicted localization of GFP and modified-GFP sequences.

We used several metrics to evaluate the reconstruction performance of CELL-E, summarized in Table S.2. Among the metrics, nucleus proportion accuracy measures how close the estimated proportion of pixel intensity within the nucleus is to the ground truth thresholded image. We believe this is the most relevant metric as it is not obscured by small spatial variations and nucleus boundaries can be obtained from the reference images. Description of other metrics and more information on the evaluation procedure can be found in Section S.3.7. Using these metrics, we performed ablations studies to optimize our model architecture and choice of protein language embedding (see Section S.2.3, Fig. S.5 and Table S.1).

While not specifically trained as a discrete localization classifier, we also performed naive comparison between CELL-E model and 1D protein localization classifiers MuLoc [19] and Subcons [20] specifically trained with annotated protein localizations. We focused on nuclear classification using a simple classification criteria on CELL-E output (see Section S.3.7), and the results are summarized in Table 1. We observed a relatively high degree of accuracy from this method compared to the task-specific models. CELL-E outperformed on the training set, and it was a close second for validation set proteins despite not seeing localization annotations during training.

**Table 1.** Nuclear Localization Prediction Accuracy.

|  | Train | Validation |
| --- | --- | --- |
| VQGAN | $0.99 \pm 0.08$ | $0.99 \pm 0.09$ |
| CELL-E | **$0.89 \pm 0.31$** | $0.72 \pm 0.45$ |
| MuLoc | $0.71 \pm 0.45$ | **$0.79 \pm 0.41$** |
| Subcons | $0.43 \pm 0.49$ | $0.69 \pm 0.46$ |

VQGAN indicates the accuracy evaluated on the ground truth threshold image passed through the VQGAN image encoder. As CELL-E selects tokens from this VQGAN to produce its outputs, these values represent the best possible performance for our model.

### 2.3 Analysis of NLS using CELL-E

As a first test to show CELL-E can recognize specific, functional sequence features, we let it predict the images for Green Fluorescent Protein (GFP), which is non-native to human, as well as GFP appended with two commonly used NLS’s KRPAATKKAGQAKKKK from nucleoplasmin [21] and PAAKRVKLD from N-Myc [22]) that drive nuclear localization of a protein. We also appended a randomly generated sequence as a control. A randomly chosen nuclear image from the OpenCell dataset was used as the morphological reference. CELL-E struggles to make any prediction with high confidence from the base GFP sequence or GFP appended with a short peptide containing a random sequence, whereas the two GFP-NLS fusions are clearly predicted to be localized within the nucleus. Therefore, CELL-E has the potential to perform computational insertion screenings for the functional sufficiency of putative localization sequence features.

Next, we examined whether CELL-E can identify NLS in a protein by computationally performing truncation/deletion studies. For this purpose, we chose DNA Topoisomerase I (TOP1), whose N-terminal intrinsically disordered region (amino acid (aa) 1-199) is essential for its nuclear localization [23]. An experimental study generated a series of deletion mutants for this region and imaged the subcellular localization in HeLa cells when fused to eGFP [24]. To computationally reproduce this study, we fed the exact sequences of the deletion mutants to CELL-E. As shown in Fig. 5, the predictions were largely consistent with the experimental data, recapturing the inability for *aa 1-67* to drive nuclear localization despite containing a putative NLS, as well as the sufficiency of *aa 148-199* as an NLS.

**Fig. 5.**
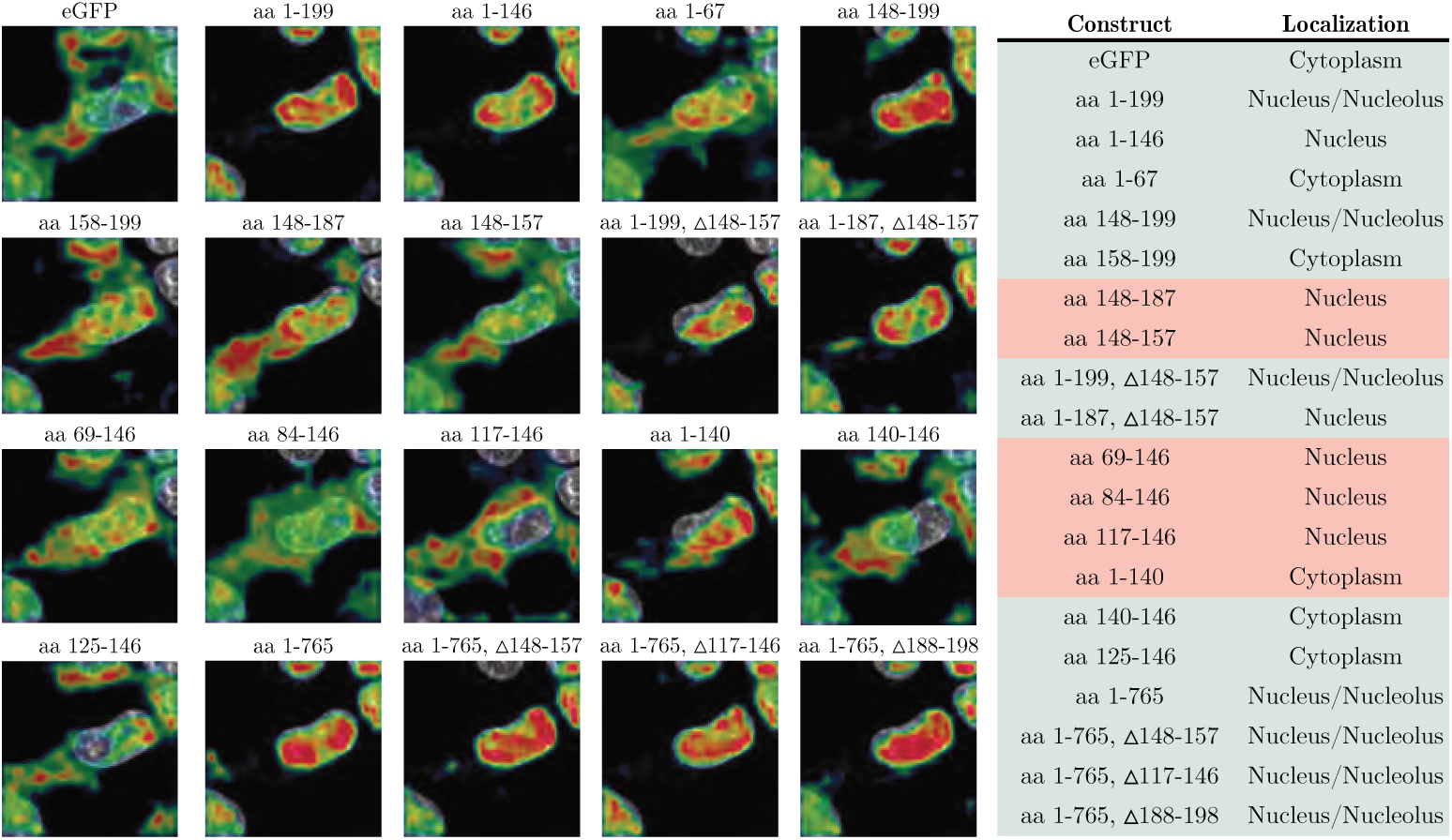
Predicted localization of eGFP fusions from [24] and corresponding figures from the original paper. *aa 1-199* contains the entire N-terminus region. *aa 1-146* only contains Motifs I and V. *aa 1-67* only contains Motif-I. *aa 148-199* contains Motif II, III, IV and V.

Lastly, we demonstrate a more direct approach than computational insertion or deletion studies to identify putative sequence features responsible for protein localization. Specifically, we split the generated image patches into two groups, one with the target protein being present and the other being absent based on the average pixel intensity within the 16 *×* 16 image patch. Then, we calculated the difference of attention weights for each amino acid token to contribute to the two groups. Fig. 6 highlights the amino acids with higher weights for the ”present” group. The highlighted amino acids includes the three putative NLSs in the experimentally verified *aa 148-199* range, as well as part of the new *aa 117-146* NLS identified in [24]. On the other hand, the putative NLS in the experimentally invalidated *aa 1-69* range are not activated. The attention map also suggest that *aa 89-107* (KIKKE) could be another NLS in this protein. We must point out that the calculation of attention map was simply based on a protein being ”present” or ”not present” in image patches and did not specify ”nuclear localization” at all. Therefore, it should be capable to serve as a general approach to discover putative sequence features driving protein localization to a variety of subcellular compartments.

**Fig. 6.**
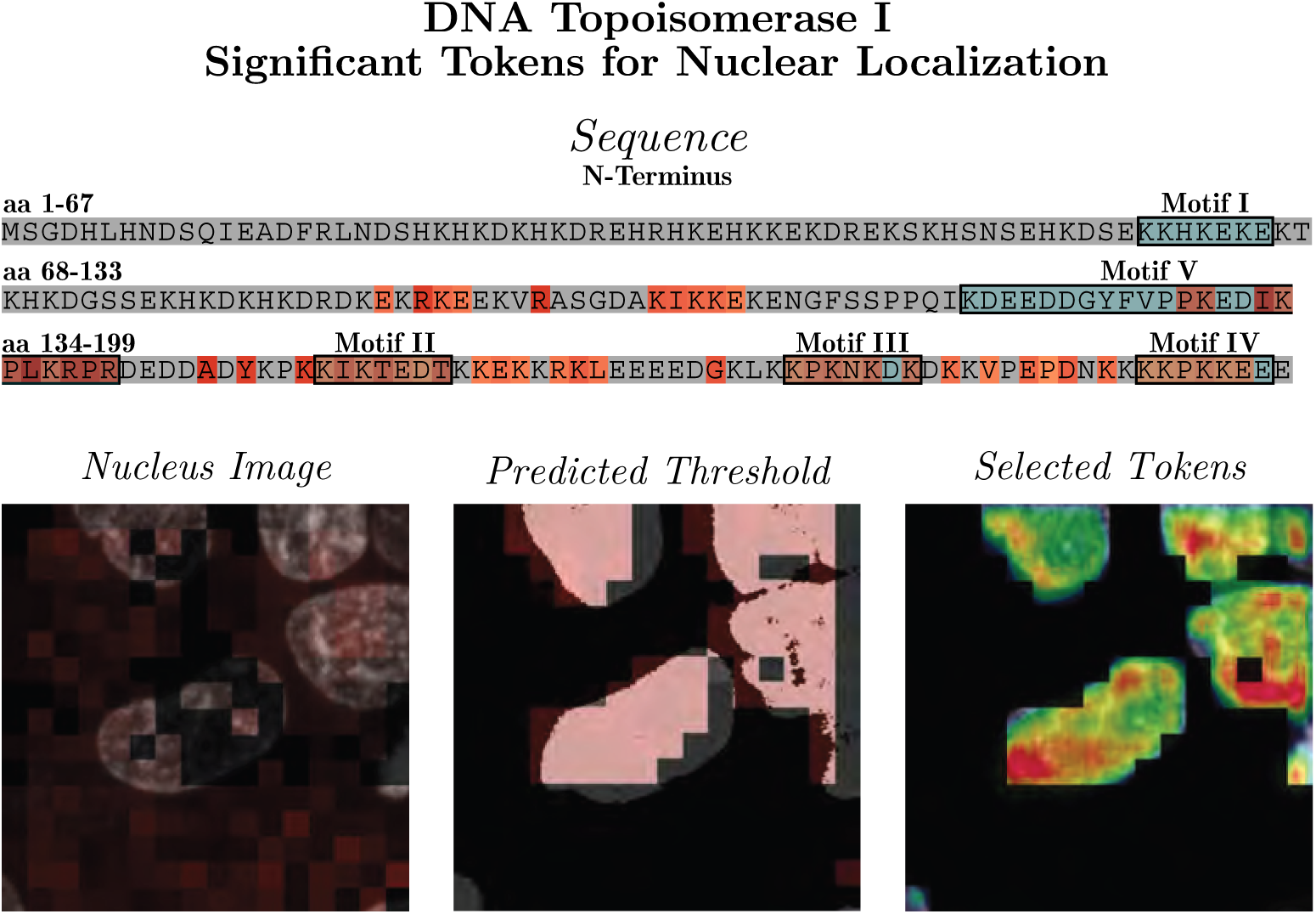
Attention weights for significant tokens when patches containing a large percentage of protein are selected (right column). Previously computationally identified putative NLSs are boxed in blue (left column). These are aa 59-65 (Motif I, **KKHKEKE**), aa 150–156 (Motif II, **KKIKTED**), aa 174–180 (Motif III, **KKPKNKD**), and aa 192–198 (Motif IV, **KKKPKKE**). Additionally, the new NLS identified in Mo et. al.[24], Motif V (aa 117-146 ), is highlighted.

## 3 Discussion

CELL-E’s performance seems to be currently limited by the scope of the Open-Cell dataset, which only accounts for a handful of proteins within a single cell type and imaging modality. As the OpenCell project is an active development, we expect stronger performance as data becomes available. The availability of brightfield (e.g. phase-contrast) images as the morphological reference will also likely improve the prediction of cytoplasmic protein localization compared to using nuclei images. Furthermore, the utility of the model comes in terms of linking embedding spaces of dependent data. One could imagine follow up experiments where rather than images being the prediction, other signatures such as protein mass spec could be predicted. Additionally, other sources of information, such as structural embeddings could be incorporated to bolster CELL-E’s capabilities.

## 4 Methods

We use a multi-phase training approach similar to DALL-E, but our model also uses pre-trained language-model input embeddings for the amino acid text sequences via TAPE:

- **Phase 1** A Vector Quantized-Generative Adversarial Network (VQGAN) [18] is trained to represent a single channel 256*×*256nucleus image as a grid comprised of 16 *×* 16 image tokens (Fig. S.7), each of which could be one of 512 tokens.
- **Phase 2** A similar VQGAN is trained on images corresponding to binarized versions of protein images. These tokens represent the spatial distribution ofthe protein (Fig. S.9).
- **Phase 3** The VQGAN image tokens are concatenated to 1000 amino acid tokens for the autoregressive transformer which models a joint distribution over the amino acids, nucleus image, and protein threshold image tokens.

### 4.1 Model Specifics

The optimization problem is modelled as maximizing the evidence lower bound (ELBO) [25, 26] on a joint likelihood distribution over protein threshold images *u*, nucleus images *x*, amino acids *y*, and tokens *z* for the protein threshold image:

**Theorem 1.**

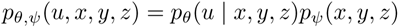

This is bounded by:

**Theorem 2.**

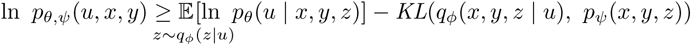

where *q_ϕ_* is the distribution 16 *×* 16 image tokens from the VQGAN corresponding to the threshold protein image *u*, *p_θ_* is the distribution over protein threshold generated by the VQGAN given the image tokens, and *p_ψ_* indicates the joint distribution over the amino acid, nucleus, and protein threshold tokens within the transformer.

### 4.2 Nucleus Image Encoder

Training both image VQGANs maximizes ELBO with respect to *ϕ* and *θ*. The VQGAN improves upon existing quantized autoencoders by introducing a learned discriminator borrowed from GAN architectures [18]. The Nucleus Image Encoder is a VQGAN which represents 256 *×* 256 nucleus reference images as 256 16 *×* 16 image patches. The VQGAN codebook size was set to *n* = 512 image patches. Further details can be found in Section S.3.4.

### 4.3 Protein Threshold Image Encoder

The protein threshold image encoder learns a dimension reduced representation of a discrete binary PDF of per-pixel protein location, represented as an image image. We adopt a VQGAN architecture identical to the Nucleus VQGAN. The VQGAN serves to approximate the total set of binarized image patches. While in theory a discrete lookup of each pixel arrangement is possible, this would require *∼* 1.16 *×* 10^77^ entries, which is computationally infeasible. Furthermore, some distributions of pixels might be so improbable that having a discrete entry would be a waste of space.

Protein images are binarized with respect to a mean-threshold, via:

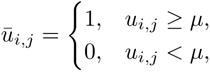

*∀* pixels *u ∈* image *U* of size *i × j*, where *µ* is the mean pixel intensity in the image (Fig. S.8).

The 16 *×* 16 image patches learned within the VQGAN codebook therefore correspond to local protein distributions. In Section 4.6, we detail how a weighted sum over these binarized image patches is used to determine a final probability density map. Hyperparameters and other training details can be found in Section S.3.5.

### 4.4 Amino Acid Embedding

For language transformers, it is necessary to learn both input embedding representations of a text vector as well as attention weights between embeddings [27]. In practice, this creates a need for very large datasets [28]. The OpenCell dataset contains 1,311 proteins, while the human body is estimated to contain upwards of 80,000 unique proteins [29]. It is unlikely that such a small slice could account for the large degrees of variability found in nature.

In order to overcome this obstacle, we opted for a transfer learning strategy, where fixed amino acid embeddings from a pretrained language model exposed to a much larger dataset were utilized. We found the strongest performance came from TAPE embeddings [16]. Utilizing pretrained embeddings had the two-fold benefit of giving our model a larger degree of protein sequence context, as well as reducing the number of trained model parameters, which allowed us to scale the depth of our network.

We tried training using random initialization for amino acid embeddings (See Section S.2.3), however, we noted overfitting on the validation set image reconstruction and high loss on validation sequences. We also experimented with other types of protein embeddings, including UniRep [30] and ESM1-b [31].

### 4.5 CELL-E Transformer

The transformer (*p_ϕ_*) utilizes an input comprised of amino acid tokens, a 256 *×* 256 nucleus image crop, and the 256 *×* 256 corresponding protein image threshold crop. In this phase, *ϕ* and *θ* are fixed, and a prior over all tokens is learned by maximizing ELBO with respect to *ϕ*. It is a decoder-only model [32].

The model is trained on a concatenated sequence of text tokens, nucleus image tokens, and protein threshold image tokens, in order. Within the CELL-E transformer, image token embeddings were cast into the same dimensionality as the language model embedding to in order to maintain the larger protein context information, however the embeddings corresponding to the image tokens within this dimension are learned (See Fig. 2).

### 4.6 Probability Density Maps

When generating images, the model is provided with the amino acid sequence and nucleus image. The transformer autoregressively predicts the protein-threshold image. In order to select a token, the model outputs logits which contain probability values corresponding to the codebook identity of the next token. The image patch *v_i_* is selected by filtering for the top 25% of tokens and applying top-k sampling with gumbel noise [33].

Ordinarily, the final image is generated by converting the predicted codebook indices of the protein threshold image to the VQGANs decoder. However, to generate the probability density map *v̄*, we include the full range of probability values corresponding to image patches, *p*(*v_i_*), obtained from the output logits. The values are clipped between 0 and 1 and multiplied by the embedding weights within the VQGAN’s decoder, *w_i_*:

**Theorem 3.**

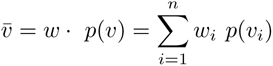

This output is normalized and displayed as a heatmap (Fig. 7).

**Fig. 7.**
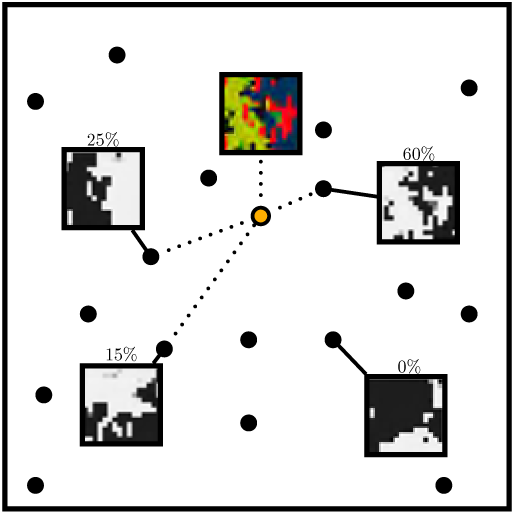
Simplified example of probability map calculation. Each circle corresponds to an image token within the quantized VQGAN embedding space. Each PDF patch (yellow) is obtained as a weighted sum over all protein threshold image VQGAN codebook vectors.

## 5 Data and Code availability

Our model is a heavily modified version of an open source text-to-image transformer [34], available via the MIT license (Copyright (c) 2021 Phil Wang). Our code is available at https://github.com/BoHuangLab/Protein-Localization-Transformer via the MIT license (Copyright (c) 2022 Emaad Khwaja, Yun Song, & Bo Huang).

## 6 Acknowledgements

B.H. is supported by the National Institutes of Health (R01GM131641). Y.S.S. and B.H. are Chan Zuckerberg Biohub - San Francisco Investigators. Y.S.S. is supported by NIH grant R35-GM134922.

## 7 Author information

E.K. played a key role in the advancement of the approach, carrying out the majority of the coding, designing and conducting a significant number of the experiments, and producing an initial version of the manuscript. The remaining authors also offered consistent input on all aspects of the project, assessed the code, and helped with the final draft of the manuscript.

## Appendix S.1 Supplementary Figures

## Appendix S.2 Supplementary Notes

### S.2.1 Related work

Natural language processing (NLP) has found applications in amino acid sequence encoding, due to the long contextual dependencies of amino acids in a protein’s folded three-dimensional structure [35]. Self-supervised models from the NLP field have demonstrated excellent performance in predicting protein properties from amino acid sequence inputs [31, 36–38]. These models are trained on millions of amino acid sequences from databases such as BFD [39], UniRef [40], Pfam [15], and Protein Data Bank [41]. The language models have proven effective in downstream tasks like structure prediction, evolutionary analysis, and protein engineering [16], with LSTM and attention-based models achieving particularly impressive results [27]. UniRep is an LSTM model that predicts the next amino acid in a variable length sequence [42], while BERT uses bidirectional masked language modeling to predict the identity of masked tokens throughout the sequence [43]. Facebook’s Evolutionary Scale Model (ESM) is a state-of-the-art masked-language model model, pre-trained with 250 million amino acid sequences and over 700 million parameters [31].

While traditional supervised approaches, such as stochastic modeling, have been limited by feature representation or computation time, deep learning has proven to be a powerful tool in predicting localization [44]. With the ability to optimize millions of parameters, deep neural networks have shown the ability to represent complex patterns in a manner that traditional manual feature extraction cannot [45, 46].The success of language models in protein prediction tasks suggests that patterns dictating these structures are buried within residue sequences [16, 30].

Protein localization is typically framed as a class prediction task. 1D localization predictors take the primary sequence as input and produce a fixed-length vector, with each entry corresponding to a subcellular location and the values being probability values. However, these methods have limitations [47, 48]. Discrete classifications for contiguous regions of the cell, such as the nuclear membrane, can be ambiguous and may have flawed annotations in established datasets. [49]. Additionally, these methods do not account for the influence of local cellular geometries [50–53] and cell states [54, 55] on transport dynamics. For example, one would expect significantly high amounts of transcription factors for DNA replication in the S-phase of the cell cycle, but not during cell separation in mitosis [56]).

### S.2.2 Text-To-Image Generation

Ramesh et. al. [10] demonstrated true zero-shot text-to-image generation with their model, DALL-E. Unlike previous models, DALL-E utilized an autoregressive framework, which was trained on a joint distribution of text and image tokens, enabling it to make novel image predictions with high fidelity. In contrast, earlier models based on variational autoencoders (VAE) [25] or Generative Adversarial Networks (GAN) [57] performed poorly when generating images outside of the training data, resulting in distorted images and artifacts [58–60].

While our method does similarly allow for truly zero-shot protein image prediction (Fig. S.3), our goal for image generation extends beyond visual fidelity and includes a degree of spatial accuracy. This is crucial for capturing the dynamic process of protein abundance in cells, which fluctuates with respect to cell state and environmental factors. To overcome this challenge, the model is tasked with predicting per-pixel binary probability representations of protein localization, which can then be linearly combined to generate a continuous 2D probability density function of protein localization.

The following images (Fig. S.3, Fig. S.4) are from text-to-image models architecturally similar to CELL-E, but replace the Protein Threshold VQGAN with a similarly trained Protein Image VQGAN.

Fig. S.4 shows model outputs model similar to Fig. S.3 above, but does include image synthesis conditioned on a nucleus image input (via Nucleus Image VQGAN). The predicted outputs are perceptually more similar to the ground truth protein image, but the questions of accuracy and scientific utility mentioned previously continue to be a factor within this paradigm.

The sequence-to-image transformer is similar to CELL-E and does not take a nucleus image as input. This produces perceptually similar images (Fig. S.3). Within the training data, the corresponding protein image is merely a snapshot in time and could be markedly different if taken at another time point. For this reason, we do not believe a image prediction without confidence provides scientific utility.

### S.2.3 Ablation Study

#### Model Architecture

In order to gauge the importance of each component of the model, we performed ablation by training several versions of the model with the same initialization. We specifically chose to look at performance in nuclear proportion accuracy over the validation set (Fig. S.5).

On the top row, performance from the VQGAN is used as a reference of possible top performance, just as in Table S.2. The second row depicts our main CELL-E (with depth = 32 and a fixed language embedding) used for this study. We found the performance in both cross-entropy and nuclear proportion accuracy increased with model depth when compared to similar models of smaller depth (third, fourth, and fifth rows).

To understand the effect of using fixed language models, we trained 2 versions of CELL-E of same depth. The first (second row, light blue) had a fixed language embedding, while the second (seventh row, red) was free to change during training. We also introduced a model with a randomly initialized language embedding (sixth row, light green). While we note fairly high performance from the unfixed models on the training data, they performed quite poorly on the validation, indicating severe overfitting. This is a result of the comparatively small number of proteins represented within the OpenCell dataset when compared to the large pFam database used to train TAPE.

We also trained a versions of the model which did not use a nucleus input (second to last row, pink) like DALL-E, and a model that only used a nucleus input and no sequence (last row, purple), although a start token was still prepended.

Overall, we observe a distributional shift to the right, indicating more accurate predictions, as the depth of the transformer is scaled. Full results for both training and validation sets can be seen in Table S.1. We also evaluated the performance of CELL-E using different protein embedding spaces. These were configured such that they were either at the same depth as the TAPE model, or the depth was scaled as deep as possible such that the GPU memory was saturated during training. All model were trained until convergence on the validation set.

#### Language Embedding

Alongside language embeddings, we also used one-hot and amino acid chemical descriptors as embedding features. The amino acid chemical descriptors come from Osorio et. al. [61], which contains amino numerical descriptions of amino acid properties from various literature sources [62–71]. Using one-hot and amino acid descriptors allowed us to scale to deeper model depths, but we did not see much improvement from doing so using these embeddings. These encodings likely do not contain sufficient information complexity about local environments that TAPE, UniRep, and ESM1b contain. While we do not see a consistent top performer on the training data, TAPE-based models generally performed the best across all metrics on the validation set, indicating a higher degree of generalizability.

## Appendix S.3 Supplementary Methods

### S.3.1 Dataset

Each protein entry in OpenCell is accompanied by multiple high-resolution 3D confocal images containing multiple cells [13]. Having multiple live cells enables the potential for protein distribution to be captured at several time points within a cell’s lifetime. To reduce computational cost for out demonstration, we converted a 3D z-stack into a 2D maximum intensity projection [72], which still clearly depicts most subcellular structures and allow subtle subcellular protein localization differences to be distinguished from the OpenCell images [13].

The OpenCell dataset was selected because the split-fluorescent protein fusion system allows for tagging endogenous genomic proteins, maintaining local genomic context, and the preservation of native expression regulation [13]. This last point is specifically important when compared to the previously mentioned HPA, which contains *∼* 10*×* more proteins and images. ICC-IF, which is the technique used for obtaining HPA images, requires several rounds of fixation and washing [73]. This means the proteins are not observed in a live cell, are subject to signal loss, artifacts, and/or relocalization events, and therefore does not represent the true nature of protein expression and distribution within a cell [74].

Training and validation sets were generated by randomly splitting the OpenCell dataset by protein 80%-20% training-validation For every stage of training, models were blind to sequences, nuclei, and protein images contained within the validation set. We utilize data augmentation techniques such as random horizontal and vertical flips on images during training.

### S.3.2 Train-Validation Split Sequence Diversity

In machine learning applications which utilize amino acid sequence, it is recommended to cluster proteins based on similarity in order to create a distributional shift between a training and validation (and/or test) set. Oftentimes, redundancy in subsequences between both sets may results in memorization of training sequences and inflated performance metrics [75].

To investigate the effect of this on CELL-E, we performed a clustered split using a procedure identical to the one used by [76] to create, a standard dataset used in benchmarking protein localization prediction. This model relies on PSI-CD-HIT [77]. In short, we clustered proteins based on a value cutoff of a designated percentage of identity for which the alignment must cover 80% of shorter sequences. We retrained CELL-E with train/validation splits with clustered with varied threshold percentages of sequence identity, ranging from 15% to 95% for 130 epochs. Our random split used for the main CELL-E effectively represents clustering based on 100% identity.

We did not observe any patterns in cross-entropy loss during training of the main transformer model in response to different cutoff values for sequence identity.

### S.3.3 Training

We utilized 4*×*NVIDIA RTX 3090 TURBO 24G GPUs for this study. 2 GPUs were utilized for training VQGANS via distributed training. Only a single GPU is ever used to train CELL-E models.

Our computer also contained 2*×*Intel Xeon Silver and 8*×*32768 mb 2933MHz DR*×*4 Registered ECC DDR4 RAM.

### S.3.4 Nucleus Image Encoder

VQGAN code was obtained from Esser et. al. [18], which was available via MIT license (Copyright (c) 2020 Patrick Esser and Robin Rombach and Björn Ommer).

The model was trained on random 256 *×* 256 crops of 512 *×* 512 nuclei images. Adam Optimizer was used with learning rate set to 4.5 *×* 10*^−^*^6^. The model was initially trained solely using mean-squared error reconstruction loss. After 50,000 steps, *∼* 7 epochs, the discriminator loss term was introduced. This terms helps with reducing the blurriness typically associated with VAEs. 512 discrete image codes were learned. Training occurred until the model reached convergence (at 344 epochs).

### S.3.5 Protein Threshold Image Encoder

The model was trained on random 256 *×* 256 crops of 512 *×* 512 nuclei images. Adam Optimizer was used with learning rate set to 4.5 *×* 10*^−^*^6^. The model was initially trained solely using mean-squared error reconstruction loss. After 50,000 steps, *∼* 7 epochs, the discriminator loss term was introduced. This terms helps with reducing the blurriness typically associated with VAEs. 512 discrete image codes were learned. Training occurred until the model reached convergence (at 371 epochs).

### S.3.6 CELL-E Transformer

Amino acid sequences were converted to indices via the selected language tokenizers. Unless otherwise stated, all results in this work utilized the IUPAC tokens and TAPE language embeddings. CELL-E uses encodings from TAPE, There are 30 possible codebook values for amino acids within this model, with 25 corresponding to amino acids and 5 corresponding to special tokens (i.e. padding). amino acid sequence length was limited to 1000 amino acids, which is longer than 96% of sequences within the dataset. For amino acid sequences shorter than 1000 amino acids, an end token (if utilized by the language model) was appended, followed by padding tokens. For amino acid sequences longer than 1000 amino acids, we randomly cropped a 1000 length subsection. If the right end of the crop ended before the true end of the amino acid sequence, no end token was applied. A start token is then prepended to all 1000 length sequence. The TAPE model used represents input embeddings as vectors with dimension *n×* 768, where *n* is the number of amino acids. The sequence embedding for the TAPE based models therefore had embedding vector sizes of 1001 *×* 768. Input amino acids were tokenized and their embeddings were retrieved from the language models. This input embedding is fixed. We also explored other embeddings (UniRep, ESM1b, One-hot encoding, and chemical descriptors) in Table S.1.

CELL-E was trained with an Adam optimizer with learning rate set to 3 *×* 10*^−^*^4^. The images were passed through the encoders of their respective VQGANs to obtain codebook tokens, and the final protein threshold image token is removed. We utilized data augmentation techniques including random cropping and random flips, just as was performed when training the VQGAN models. Within the CELL-E transformer, image token embeddings were cast into the same dimensionality as the language model embedding to in order to maintain the larger protein context information, however the embeddings corresponding to the image tokens within this dimension are learned. This ultimately creates a full sequence embeddings 1512 *×* 768 (1001 *×* 768 for text, 256 *×* 768 for nucleus images and 255 *×* 768 for protein threshold images) (Fig. 2).

A rotary positional embedding [78] is then applied to the input embeddings. We noted improved performance by shifting embeddings over by 1 (time-shifting [79]) in the feature dimension, but only for image tokens. Image token embeddings were shifted one position from the top and one position from the left.

A full attention scheme [80] is used where future tokens are masked in order to retain full sequence and image context. The output of the attention layers is passed through a block consisting of a linear and softmax layer to produce logits for predicted tokens at each position. Selective masking is applied so the model is unable to select anything but amino acid tokens for amino acid positions, nucleus image tokens for the nucleus image positions, and protein image tokens for the protein image positions.

Cross-entropy loss is used to measure the model’s ability to reconstruct the original input vector (without the prepended start token and including the removed final protein threshold image token). The cross entropy is initially scaled by the length of the input, but further weighting is placed to emphasize the output protein image threshold tokens. We used weightings of 1/9 for the amino acid tokens, 1/9 for nucleus image tokens, and 8/9 for the protein threshold image tokens.

The main CELL-E model had a depth of 32, indicating 32 consecutive attention and feed forward blocks, and 16 attention heads with dimension = 64. We used attention and feed forward attention dropout both = .1 during training. The language embedding was fixed. Model convergence occurred at 130 epochs and these weights were used for study.

### S.3.7 Performance Evaluation

To assess performance, we generated a single prediction per image found in the Open-Cell set. Each image was randomly cropped and flipped similar to training, but cropped regions and flips were maintained between models.

#### Nucleus Proportion Accuracy

To calculate the proportion of intensity in the nucleus, we first create a mask (Fig. S.10 of the nucleus channel using Cellpose [81]. We take a sum over the predicted 2D PDF pixels found within the nucleus mask, and divide this by the sum of pixels across the image.

For the the ground truth, we use a similar masking calculation, but consider the values of the ground truth protein image. These values are subtracted to calculate a mean-average error (MAE). Since the maximum possible value is 1 and minimum possible value is 0, we report accuracy as 1 - MAE.

#### Predicted Threshold Pixel Accuracy

We simple calculate a pixel-wise MAE between the predicted protein threshold image of CELL-E and the ground truth protein threshold image.

#### Predicted 2D PDF Pixel Accuracy

This metric is similar to Predicted Threshold Pixel Accuracy, except we evaluate the difference using the predicted 2D PDF, rather than the predicted protein threshold image. We expect this number to be less accurate as tokens with less confidence will reduce the pixel value, while all values in the protein threshold image are 0 or 1.

#### SSIM

Structural similarity index measure (SSIM) is a measure of local perceptual similarity between images. It considers neighboring pixels to evaluate loss contextually by incorporating luminance and contrast information. SSIM values range between 0, indicating no similarity, and 1, indicating maximum similarity.

#### IS

Inception score (IS) is often used to evaluate the image outputs of GANs as a measure of “realisticness.” It rewards image variety and similarity to real-life data. Performance evaluation is based on the magnitude of the IS score.

#### FID

Fréchet Inception Distance (FID) is another popular metric for evaluating the quality of images from generative models. It compares the distributions between generated and ground truth images as the squared Wasserstein metric between two multidimensional Gaussian distributions. For this study FID was scored against the training or validation sets when applicable, rather than the entire OpenCell dataset.

#### Nuclear Localization Prediction

The ground truth label for nuclear localization was designated by masking the nucleus, but computing the proportion of intensity on the ground truth thresholded protein image. If *>* 50% of this intensity was contained within the area of the nuclear mask, the assigned label would be positive for nuclear localization. Otherwise, the protein would be designated as non-nuclear. For the predicted label, we took a summation over the masked and unmasked regions of the predicted 2D PDF. If *>* 50% of pixel intensity for the 2D PDF was found in the nucleus, it was classified as a nuclear localizing protein. The protein localization prediction models were provided the amino acid sequence and were considered to predict nuclear if present in the localization prediction. These predictions were also compared against our näıve labels.

### S.3.8 Visualizing Attention

To obtain Fig. 6, we first split the 16x16 generated threshold image patches into 2 groups, one where protein is primarily determined to be present *ū_i,j_ > .*75 and another where background tokens are primarily selected *ū_i,j_ < .*25. For each respective group, we calculate a median of the attention matrices and used attention rollout [82] to recursively multiply across 32 layers. The final layers of both groups are then compared. We initially look at tokens with higher weightings for the present protein group, and discard the rest.

We show the entire image generation process, with frames corresponding to time-steps, in the attached video file: DNAtopoisomerase1.mp4.

---

**Fig. S.1.**
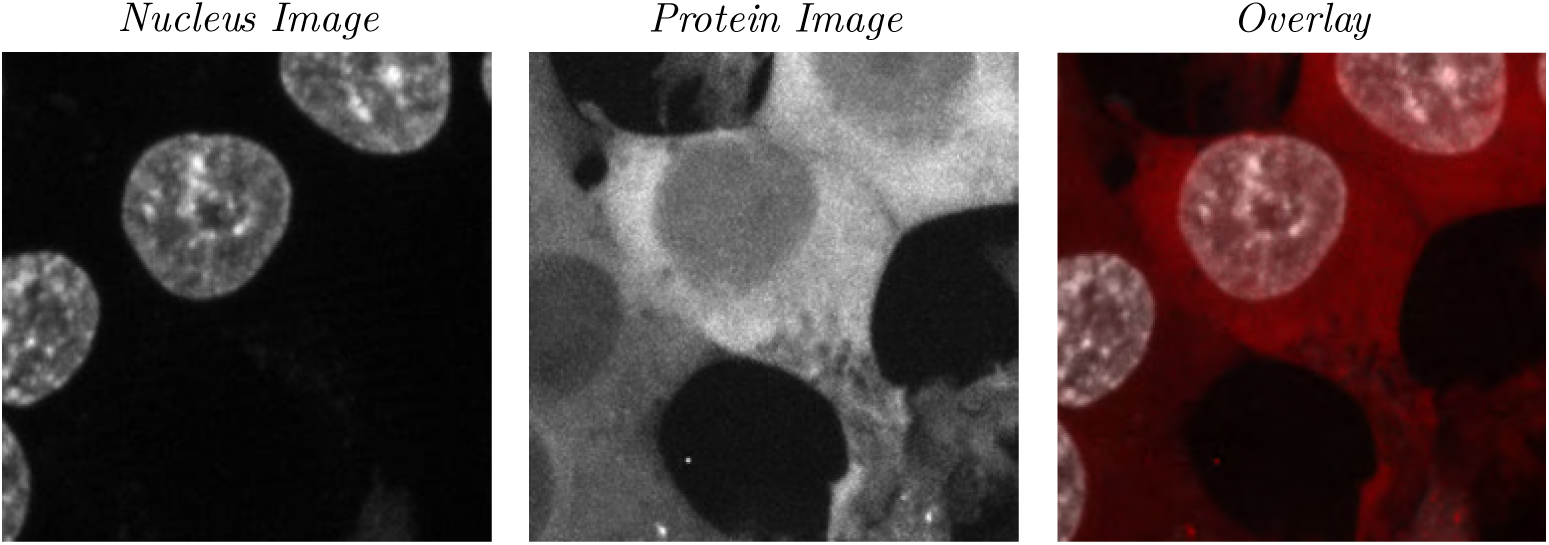
Nucleus Image (left), Protein Image (middle), and Overlay (Right). The alpha value for the protein channel in the right column is set to .7. Overlay is used as the “Original Image” in Fig. 3 and Fig. S.2.

**Fig. S.2.**
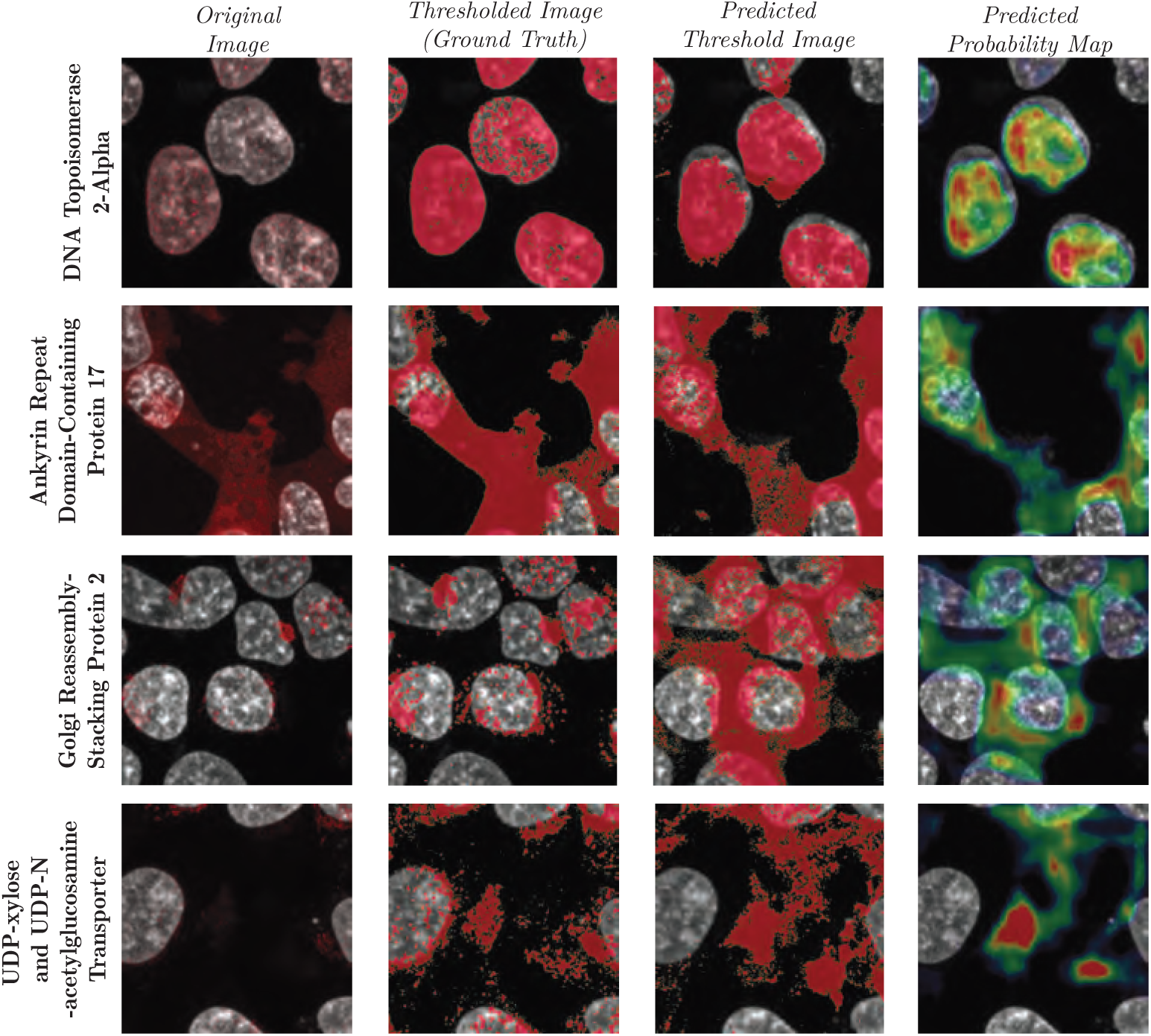
More prediction results from the validation set. We observe a high degree of spatial awareness from the model, notably in UDP-xylose-acetylglucosamine Transporter, which accurately predicts signal between cell nuclei with high confidence.

**Fig. S.3.**
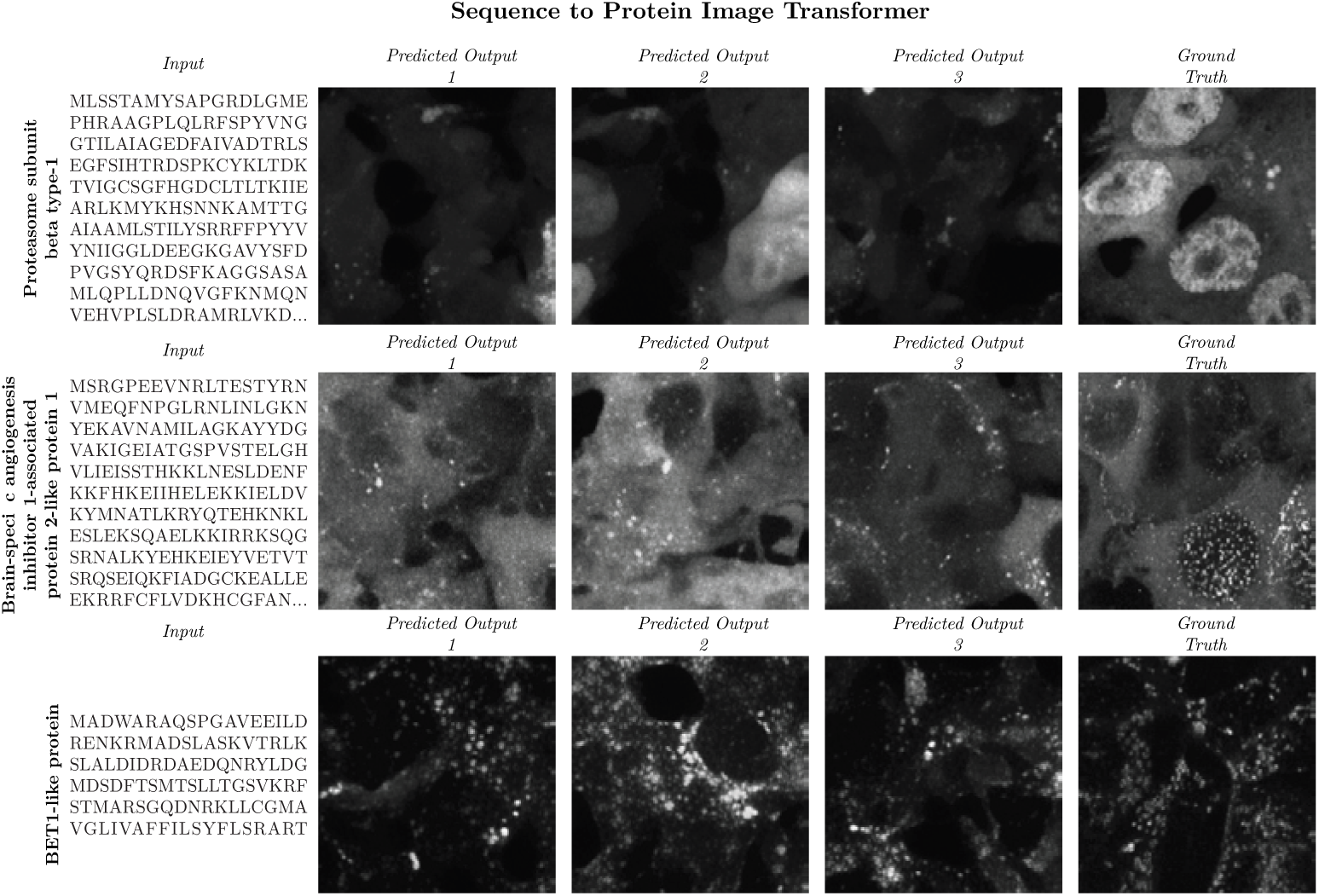
DALL-E-like model with only amino acid sequence as the input. The sequence (left column) is used as input. The middle 3 columns show separate predicted images from random initialization. The true protein image is shown in the right column.

**Fig. S.4.**
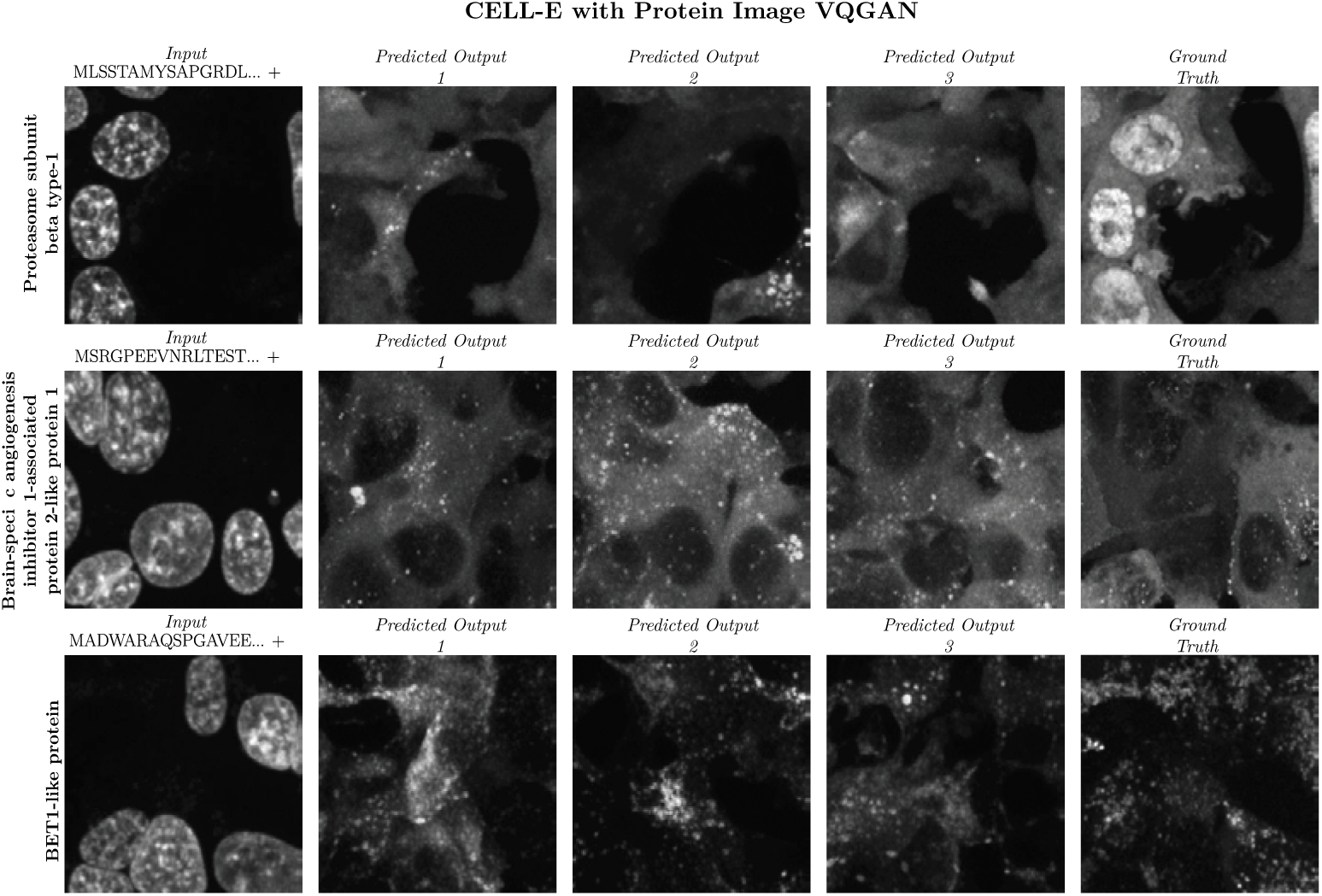
CELL-E model with Protein Threshold VQGAN replaced with Protein Image VQGAN.

**Fig. S.5.**
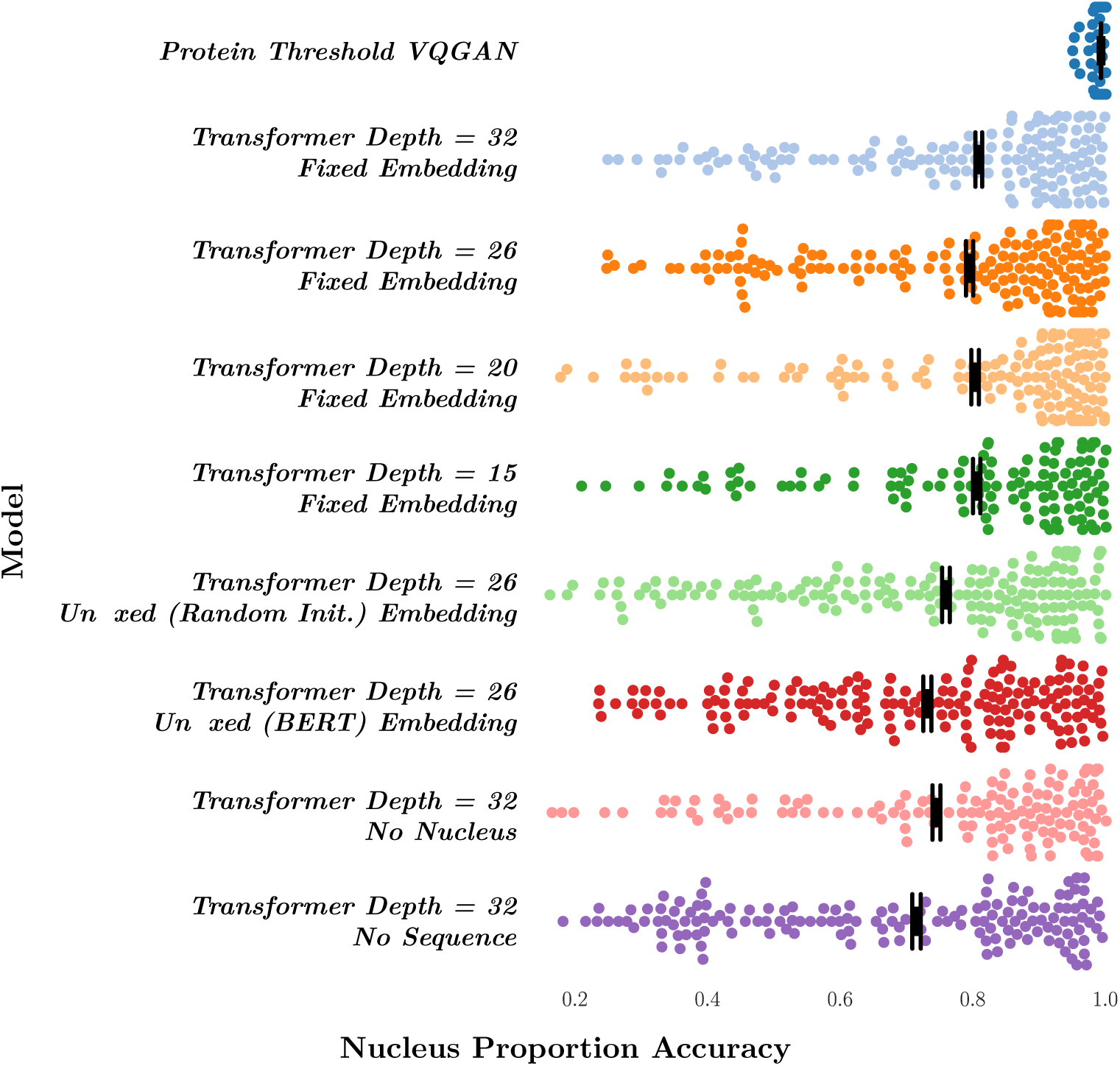
Ablation Plot. “Fixed” and “Unfixed” embedding refer exclusively to the amino acid embeddings. Image embeddings are always unfixed. Mean values and standard deviation are marked in black. *∼* 150 points are randomly selected for display out of 1303 total predictions per model.

**Fig. S.6.**
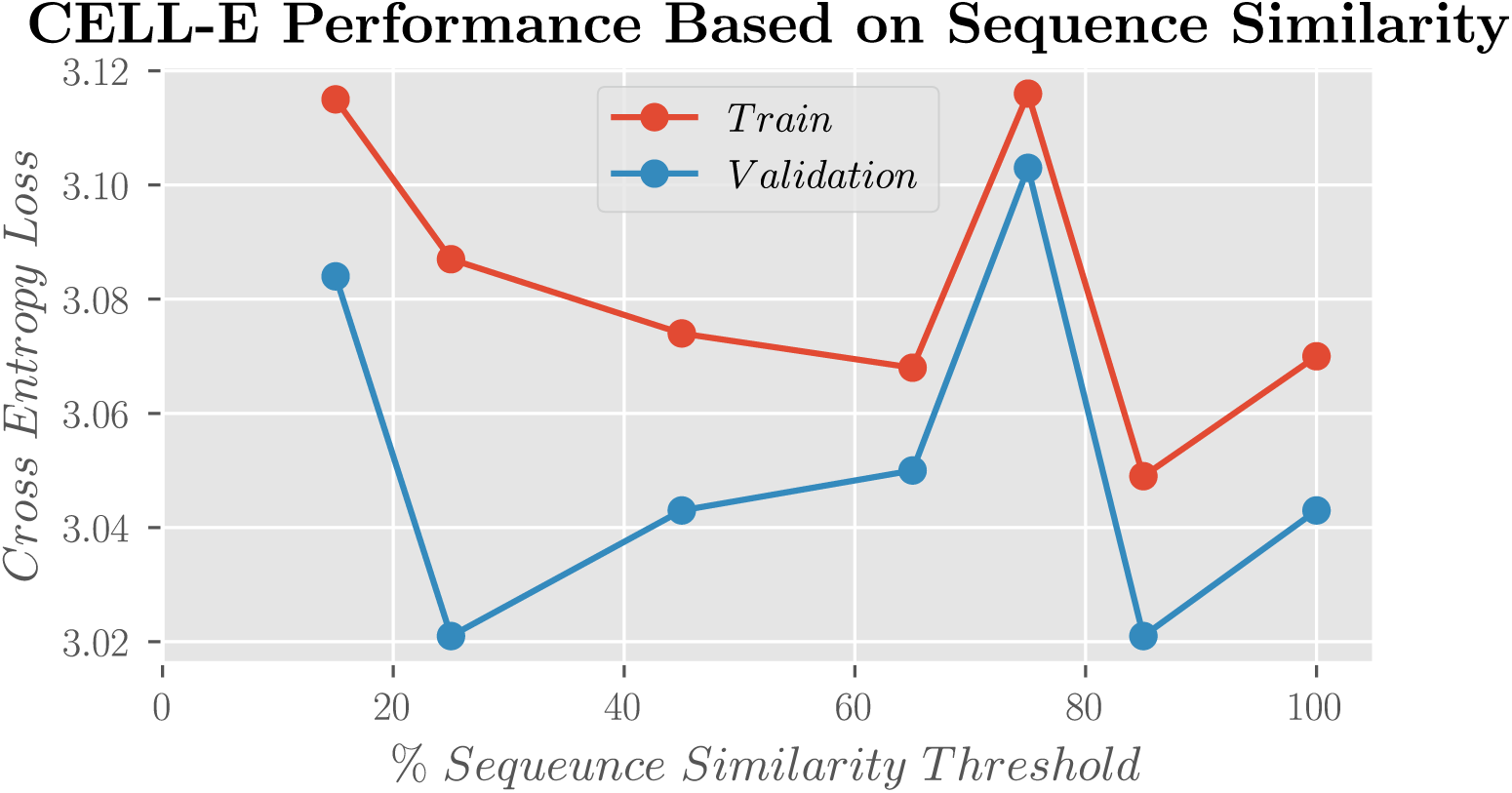

**Fig. S.7.**
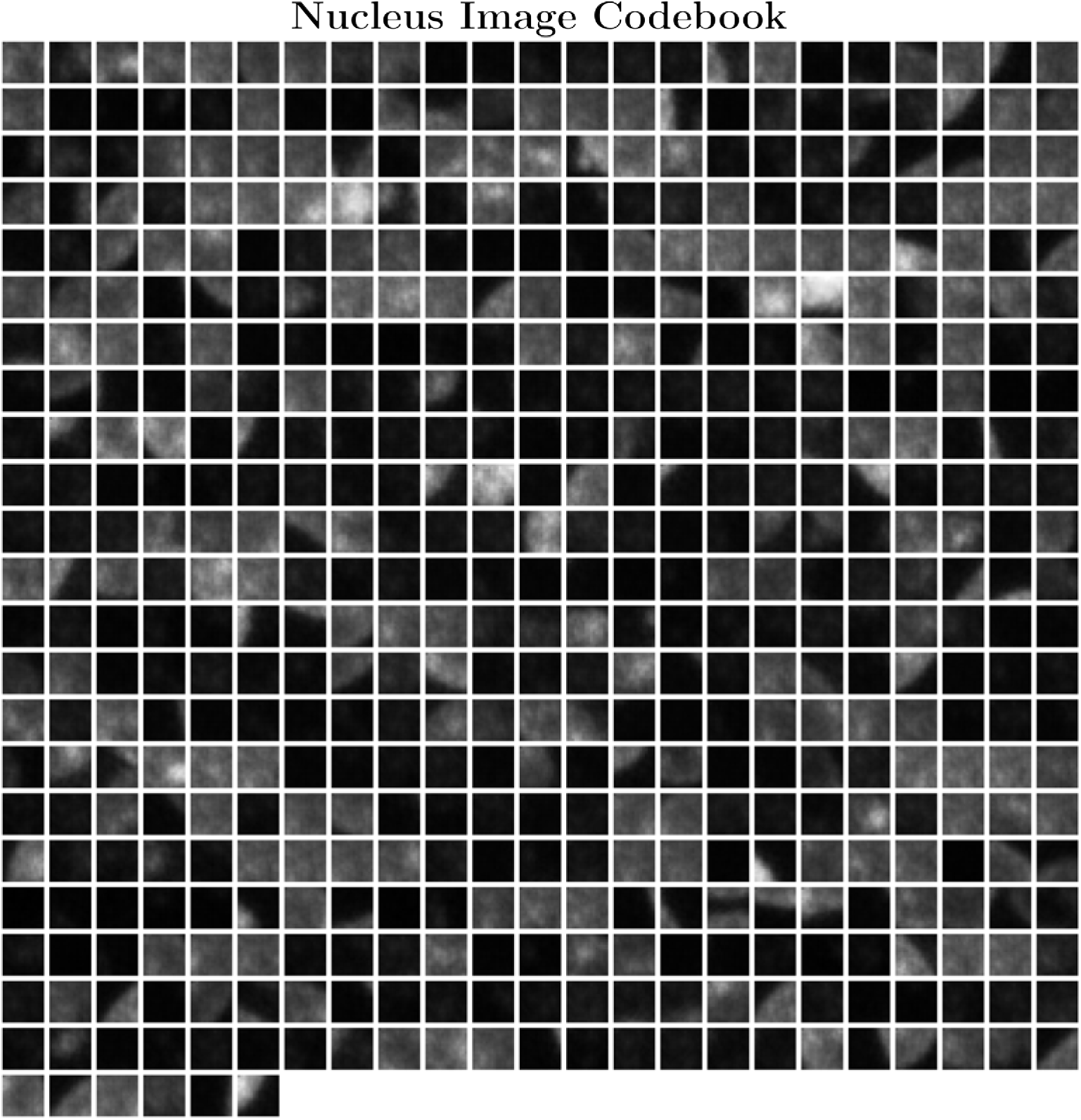
512 image patches extracted from the nucleus reference VQGAN

**Fig. S.8.**
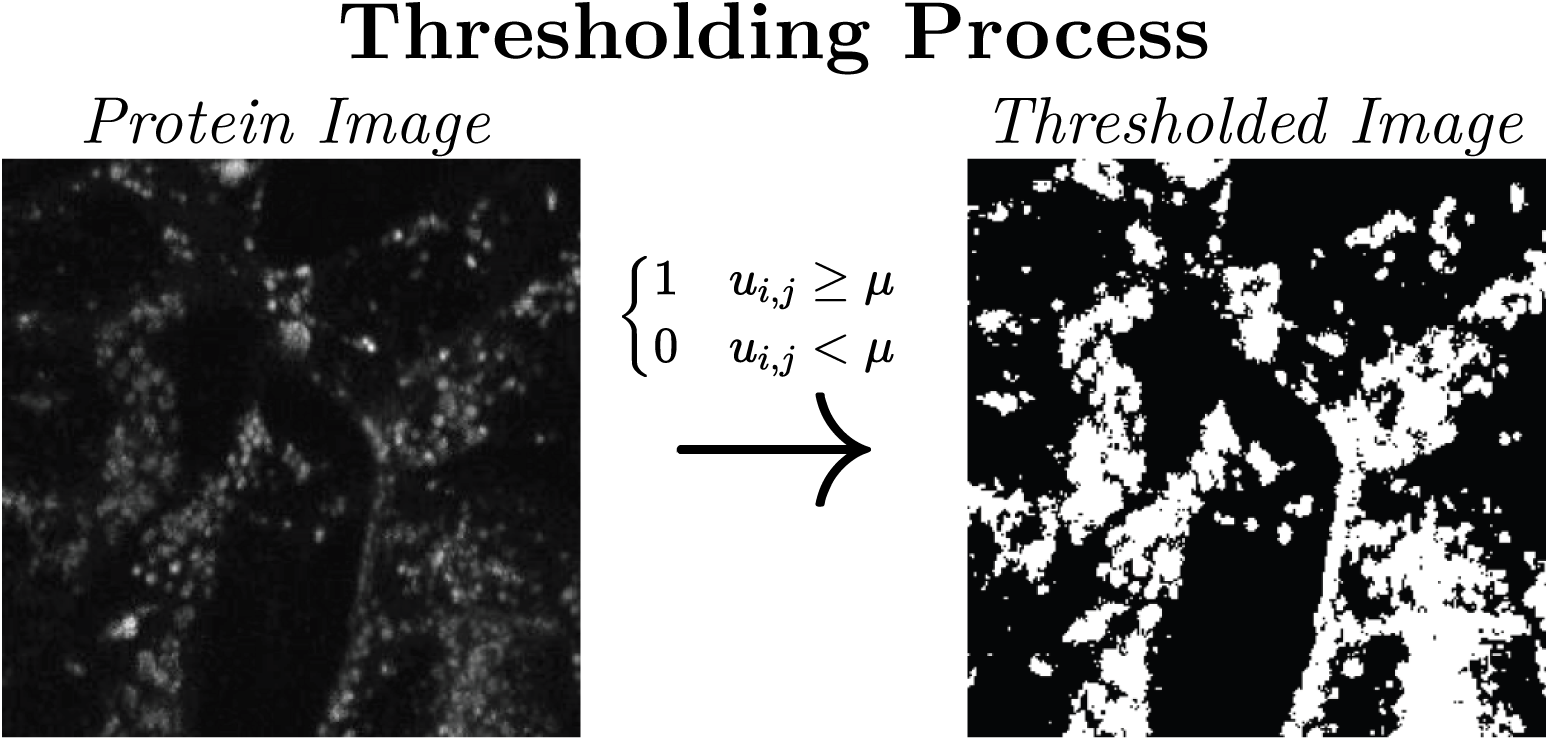
Example of thresholding process to convert protein image (left) to thresholded ground truth image (right) for CELL-E model

**Fig. S.9.**
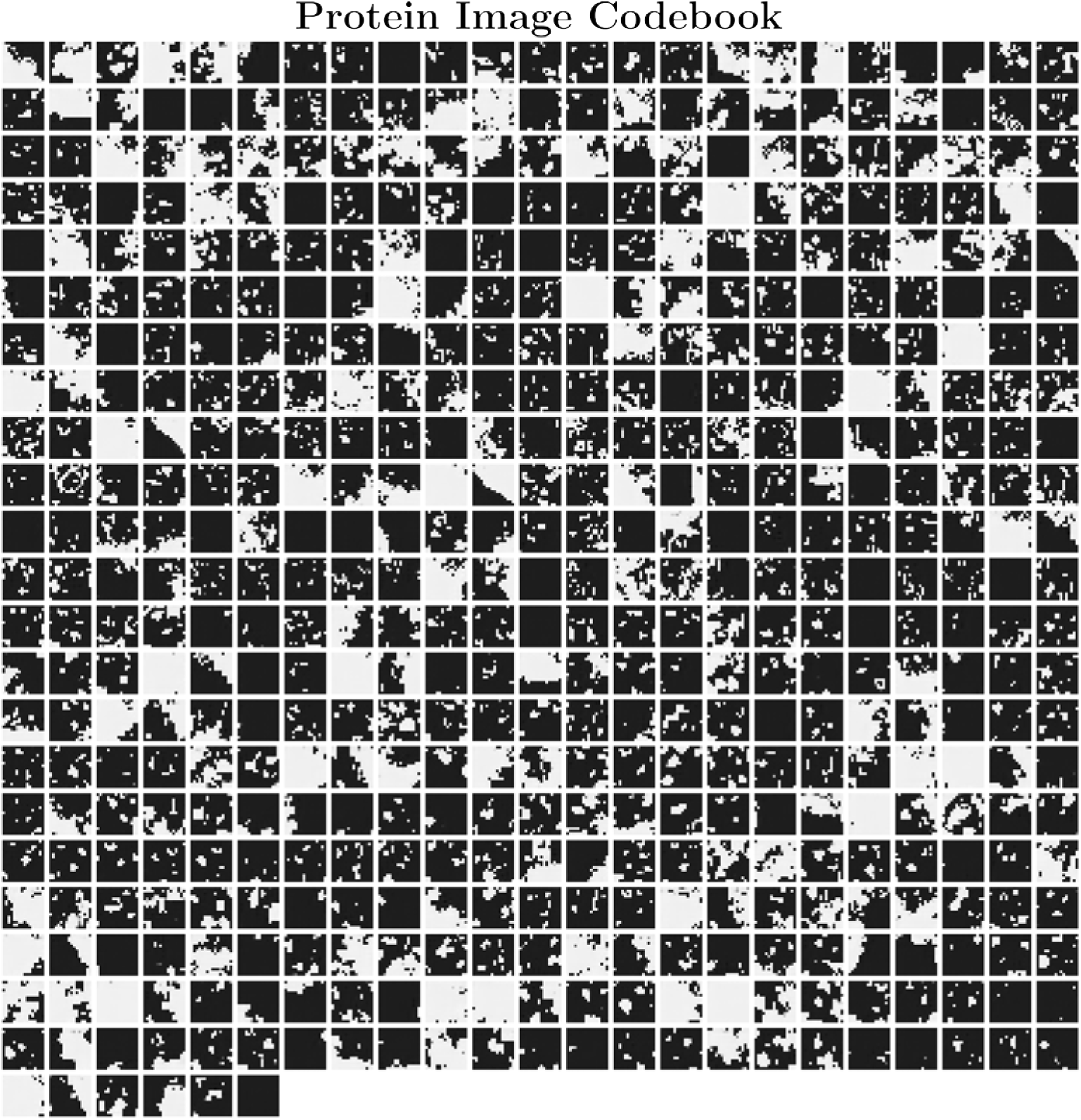
512 image patches extracted from the protein threshold VQGAN

**Fig. S.10.**
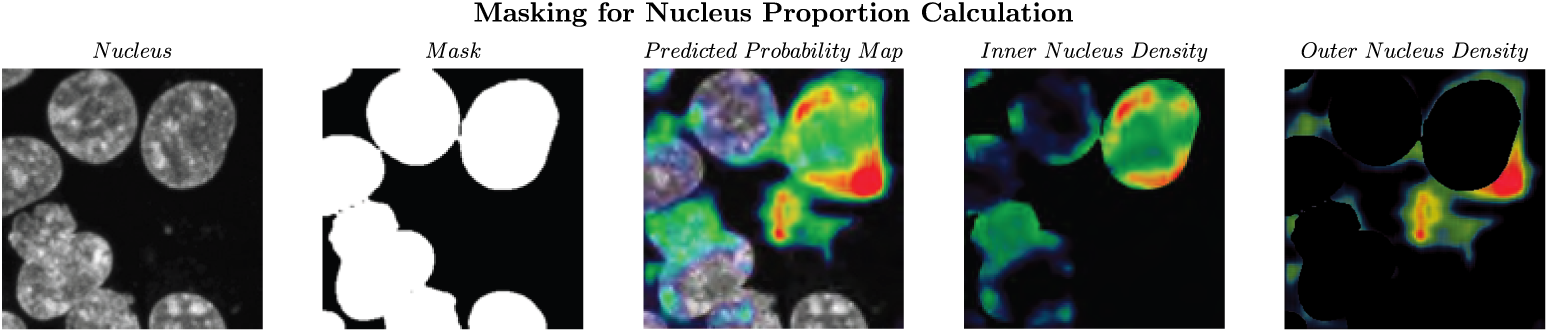
Masking procedure depicted.

**Table S.1.**
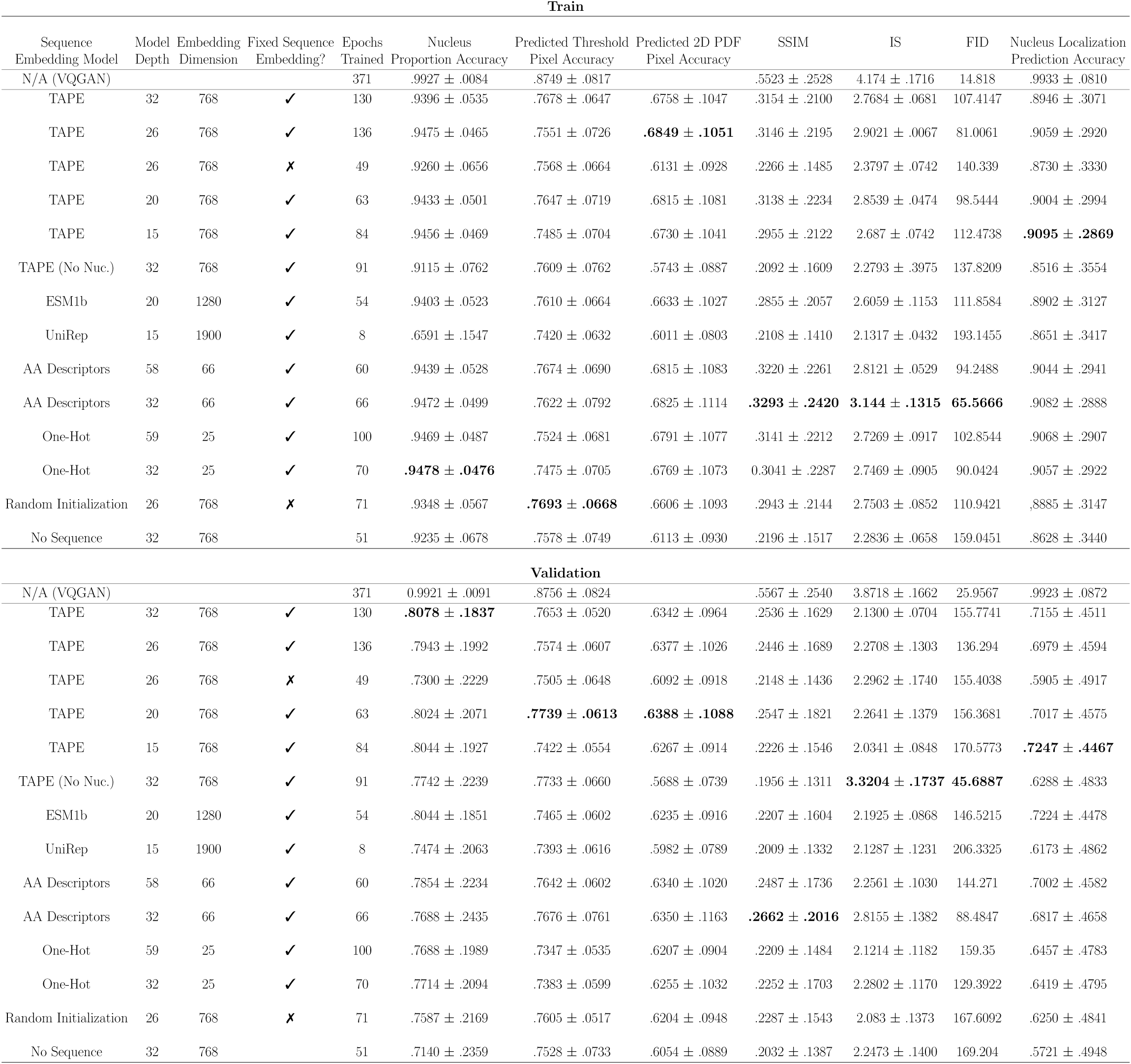
Full Results Table.

**Table S.2.** Image Accuracy.

|  | Train |  | Validation |  |
| --- | --- | --- | --- | --- |
|  | CELL-E | VQGAN | CELL-E | VQGAN |
| Nucleus Proportion Accuracy | $0.94 \pm 0.05$ | $0.99 \pm 0.01$ | $0.81 \pm 0.18$ | $0.99 \pm 0.01$ |
| Predicted Threshold Pixel Accuracy | $0.77 \pm 0.06$ | $0.87 \pm 0.08$ | $0.77 \pm 0.05$ | $0.88 \pm 0.08$ |
| Predicted 2D PDF Pixel Accuracy | $0.68 \pm 0.10$ | | $0.63 \pm 0.10$ | |
| Structual Similarity Index Measure | $0.32 \pm 0.21$ | $0.55 \pm 0.25$ | $0.25 \pm 0.16$ | $0.56 \pm 0.25$ |
| Inception Score | $2.77 \pm 0.07$ | $4.17 \pm 0.17$ | $2.13 \pm 0.07$ | $3.87 \pm 0.17$ |
| Fréchet Inception Distance | 107 | 15 | 156 | 23 |

Performance is reported as mean ± standard deviation where applicable. The VQGAN columns indicate the value of these metrics evaluated on the ground truth threshold image passed through the protein threshold VQGAN. As CELL-E selects tokens from this VQGAN to produce its outputs, these values represent the best possible performance for our model.

